# CytoScan: Automated detection of technical anomalies for cytometry quality control

**DOI:** 10.1101/2025.09.24.678276

**Authors:** Tim R. Mocking, Felix Zwolle, Yejin Park, Angèle Kelder, Yvan Saeys, Jacqueline Cloos, Sofie van Gassen, Costa Bachas

## Abstract

Studies evaluating cellular phenotypes by cytometry techniques are increasingly facing analytical challenges due to the multitudes of samples and parameters that are evaluated concurrently. Spurious technical effects resulting from a lack of standardization can affect marker distributions and further complicate multi-sample analyses. User-friendly tools for exploratory data analysis to identify such technical effects in large datasets are lacking. To fill this gap, we present a novel R package, CytoScan, that evaluates inter-measurement variation in cytometry datasets and allows for detecting anomalous measurements after data acquisition. CytoScan can detect two types of anomalies: files with limited similarity to others within a dataset (outliers) and files with limited similarity to previously acquired high-quality reference data (novelties). Using simulations of skewed marker distributions and real-life technical effects we demonstrate that CytoScan can accurately detect such anomalies. CytoScan can be applied to large cytometry datasets on consumer-grade hardware with informative visualizations, providing accessible quality-control for more reliable analyses.

## Introduction

During the past decade, advances in cytometry have enabled the high-dimensional analysis of millions of cells and their relationships within and across samples (1). However, when standardization efforts are insufficient, technical effects such as fluorochrome or antibody switches, variations in processing, or wrong instrument settings may perturb biological signals. This can result in anomalous data with skewed characteristics, leading to suboptimal or flawed analyses and potentially wrong biological conclusions.

Aside from standardization efforts to mitigate technical effects, different strategies to detect and overcome these effects in cytometry data have been explored. In cases of known technical confounders (e.g., different instruments, settings, fluorochromes), batch correction algorithms such as CytoNorm (2) and cyCombine (3) can perform inter-sample normalization. When technical effects are observed without a known cause, intra-sample normalization or scaling can transform marker expressions to the same range (e.g., 0 – 1). More complicated methods that detect and subsequently align “landmarks” in single channels across samples such as the peaks (modes) in marker distributions have also been proposed (4). However, these tools often underperform in the absence of homogeneous expression patterns or proper controls and can obscure meaningful biological signals.

When existing options are insufficient or inapplicable to overcome technical effects, removing, re-measuring or transforming only a subset of the data can be an appropriate measure to retain the biological signal in non-anomalous samples. However, in large and high-dimensional cytometry datasets, manual inspection of marker distributions to identify anomalous files is often infeasible. Consequently, anomalous samples may go unnoticed and affect biological interpretation.

Although outlier detection is a common feature of exploratory data analysis, previous tools for cytometry data analysis have primarily focused on intra-sample anomalies such as doublets (5-7), margin events (5, 8) and flow rate anomalies (5, 8, 9) rather than inter-sample anomalies. Without exploratory analyses, such anomalies would only be identified during downstream analysis when sample-specific signals appear after dimensionality reduction or unsupervised clustering. Alternatively, features from individual files can reveal global differences that may reflect batch effects or outlier samples. For example, the recently developed R package CytoMDS can generate summary visualizations based on inter-sample distances (10). However, CytoMDS loads all files into memory and calculates distances in a pairwise manner between all files. Consequentially, analysis of large datasets is infeasible without dedicated hardware. Additionally, the identification of potentially anomalous files relies on a manual selection rather than using an objective or data-driven measure. Consequently, there remains a gap for scalable and objective detection of anomalous cytometry data.

Here, we present a new R package called CytoScan that allows for a data-driven identification of anomalous files in large cytometry cohorts in a fast, intuitive, and interpretable manner. This analysis can be performed in two settings. CytoScan can identify anomalous files without prior knowledge (outlier detection) or identify files that deviate from a set of controls (novelty detection).

## Methods

### Input

CytoScan requires a list of flow cytometry standard (FCS) files as input. The input data consists of “test data” that is assessed for anomalous characteristics (outliers, novelties) (Figure 1A). If available, data that is manually selected as high quality data can be supplied as “reference data”. Reference data is not evaluated for anomalous characteristics. Beyond input FCS files, users can also add experimental metadata for integration in exploratory visualizations. A typical workflow on fluorescence parameters assumes that all data was acquired with the same panel. Although identifying inconsistencies in assumed panels is feasible with CytoScan, we recommend using the FCS metadata directly where possible.

**Figure 1.**
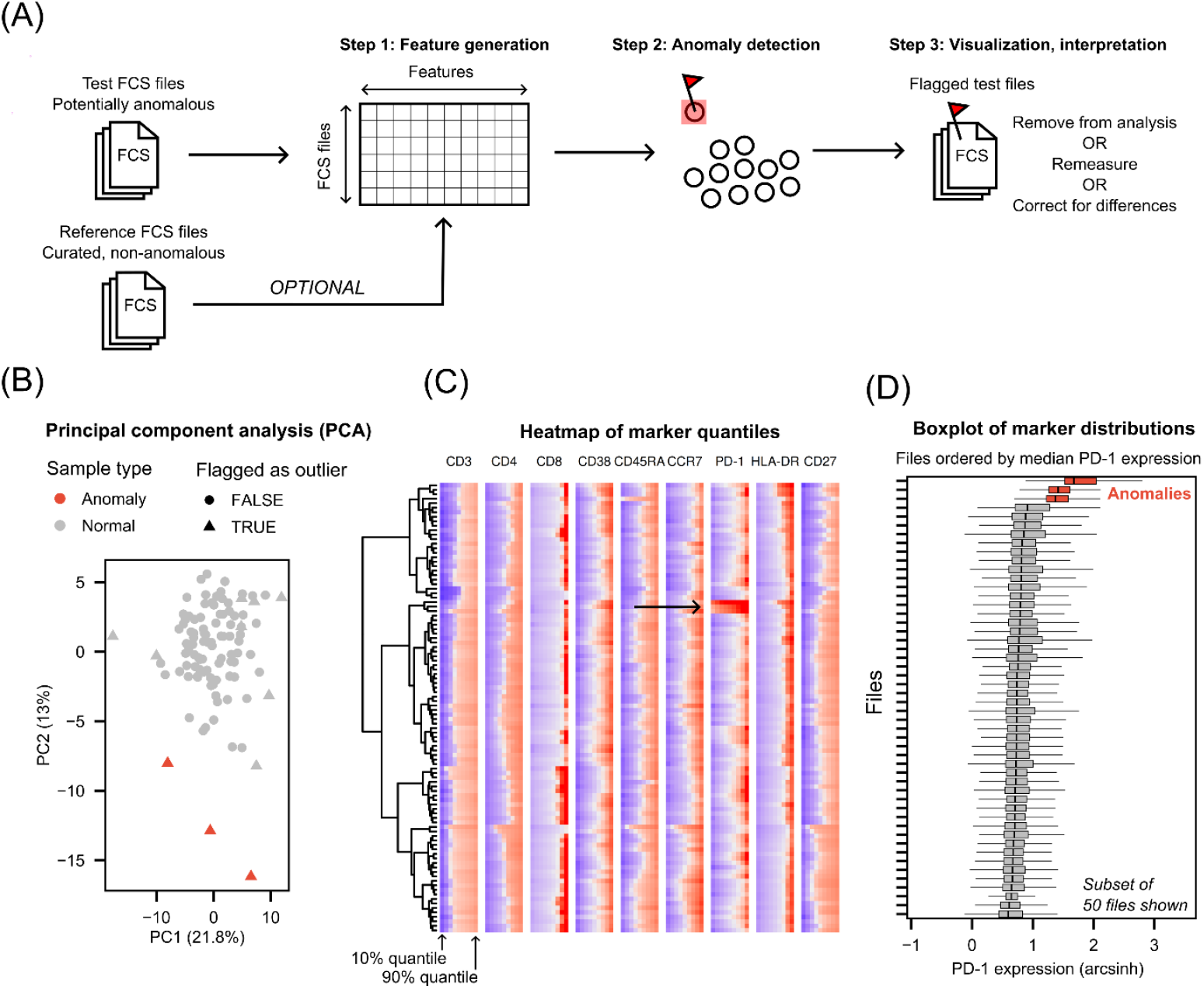
Schematic overview of CytoScan functionalities. (A) CytoScan generates features from FCS files which are subsequently used for anomaly detection using either outlier detection or novelty detection. If novelty detection is performed, a set of reference FCS files is required. Downstream visualizations support exploratory analysis and interpretation. (B-D) Examples of downstream visualizations (PCA, heatmap, boxplot) for the Liechti_28 cohort. For visualization purposes, only 10 markers are used in this example. In three random samples, we constitutively increased the signal of PD-1 on all cells to represent a file-level anomaly.

By default, CytoScan performs a minimal preprocessing by removing margin events, applying compensation (based on the $SPILL keyword in the FCS file) and transforming marker expression data using an hyperbolic arcsin transformation (co-factor 150). However, in most cases, custom pre-processing that aligns with the characteristics and objectives of the data analysis is required. This can be done by any means, as the package allows for defining a function to process each input file.

### Feature generation

CytoScan extracts features from user-defined channels in each input file. We recommend two types of features that can both be calculated for individual channels: quantiles, and the earth mover’s distance (EMD). Other features such as summary statistics (mean, median, standard deviation, interquartile-range), binning and flowFP “fingerprints” (high-dimensional binning) (11) (Table S1) are included in the package but have not been extensively tested due to inferior performance in preliminary experiments (data not shown), or challenges in downstream interpretation.

To ensure feasibility for large datasets, feature generation is performed on downsampled data. Using downsampled files still reliably captured the same global patterns as the full data (Figure S1). By reading a maximum number of randomly selected events (default: 1,000) from each file, CytoScan can generate features for large datasets on consumer-grade hardware. The user can adapt this number in case of interest in rare events. Parallelization of the feature generation process can further reduce the runtime, especially in high-performance computing (HPC) environments.

### Feature generation: quantiles

The quantile-based feature generation determines the signal intensity at several quantiles of individual channels, representing global characteristics of each channel’s distribution. To identify these features, CytoScan obtains the signal intensity by a user-defined inter-quantile distance d. By default, the 0.1 quantiles are obtained for every marker (d = 0.1). CytoScan excludes the minimum and maximum values as quantiles, resulting in 9 features per channel.

### Feature generation: earth mover’s distance

The earth mover’s distance (EMD) is a distance metric that quantifies the dissimilarity between two distributions. As comparing all possible marker distributions between all files is computationally infeasible for larger datasets, CytoScan calculates the EMD with the transport package (v0.15-4) based on a pooled marker distribution containing an equal contribution of cells from all samples. In a novelty detection setting, this pooled dataset consists of only reference samples.

### Anomaly detection

Based on the generated features, CytoScan can detect two types of anomalies: outliers and novelties. Outliers deviate significantly from other test files, while novelties are evaluated independently based on their dissimilarity to (curated) reference data. Files can be evaluated based on multiple feature generation methods and in combination with both outlier and novelty detection in the same workflow, facilitating a flexible exploratory tool for identifying anomalies.

### Anomaly detection: outliers

CytoScan uses the isotree (v0.6.1-4) implementation of the isolation forest algorithm (12) for outlier detection. This algorithm randomly selects and partitions the generated features using random splits, dividing samples in a tree structure. The algorithmic assumption is that the number of splits required to isolate a sample in this tree structure is a measure of (ab)normality. Like most decision-tree based algorithms, the isolation forest algorithm contains many hyperparameters. However, two of these are of most practical importance for CytoScan: the number of trees (*nTrees*), and the outlier score above which a cell is labeled as an outlier (*outlierCutoff*). As the isolation forest randomly selects a finite number of variables to randomly split, anomalous signals in large feature sets might not be evaluated. For this reason, increasing *nTrees* can increase the probability that a feature is evaluated by the isolation forest. As a result, anomalous signals that may be associated within one channel of a large panel might be picked up more easily.

### Anomaly detection: novelties

CytoScan uses a Gaussian mixture model (GMM) to identify novelties using a concept we described previously for cell-level anomaly detection (13). First, a GMM is fitted on the reference data using the mclust package (v6.1.1) and optimal model parameters are selected based on the goodness-of-fit defined by the Bayesian information criterion (BIC). After fitting the GMM, the log-likelihood of each reference file is calculated. Based on the distribution of log-likelihood scores, test files are labeled as novelties if the log-likelihood of that file is below a defined percentile (by default: 5%). This threshold can be adjusted: lower values result in a lower probability of detecting anomalous files (at the cost of sensitivity) while higher values increase it (at the cost of specificity).

### Visualization

In CytoScan, we implemented three types of visualizations to interpret anomalies: heatmaps, principal component analysis (PCA) and boxplots (Figure 1B-D). The exploratory nature of these plots can be further expanded by adding metadata about experimental conditions. For interpretation of plots depending on generated features, we recommend using the quantile-based features. This is because the EMD is a distance metric that can have an identical magnitude for completely opposite effects.

### Code availability

The CytoScan R package is available at https://github.com/AUMC-HEMA/CytoScan. Scripts used for performance evaluation in R are available at https://github.com/AUMC-HEMA/CytoScan-manuscript.

### Datasets

Three different cytometry cohorts (Liechti_28, BLN_BUE, HO132) were used to evaluate performance. The first dataset (Liechti_28) was selected to represent a high-quality dataset without notable technical anomalies and was used for simulating anomalies in a controlled setting. The FCS files used in this study are a random subset (n= 100) of the highly standardized 28-color immunophenotyping study (14). Samples with low T-cell viability (<25% of all files) as reported in the original study were excluded and pre-processing was performed as previously described in the original study. The second dataset (BLN_BUE) comprises two eight-color MFC cohorts (BLN Dura, BUE Dura) of acute lymphoblastic leukemia patients (ALL) (15) obtained from FlowRepository (FR-FCM-ZYVT). dPreprocessing was performed as previously described.The last dataset (HO132) comprises FCS collected during the course of the HOVON-SAKK-132 phase III clinical trial for acute myeloid leukemia patients (16). Two subsets were created focused, one focused on scatter parameters (HO132_scatter) and one focused on a specific timepoint-matched subset of measurements performed with a standardized leukemia stem cell (LSC) assay (HO132_LSC) (17, 18). Pre-processing was performed with default settings.

Involvement of human participants was approved by the applicable institutional review boards and the study followed the Declaration of Helsinki with written informed consent from all participants.

### *In silico* simulation of technical effects

In the Liechti_28 dataset, we skewed marker distributions in individual FCS files by introducing three different realistic technical effects: stretching, shifting, and separation of positive and negative peak signals. For the stretch effect, the channel was multiplied by a given effect size (1 – 2, steps of 0.05). For shifting and separation, the final effect size was determined based on the interquartile range (IQR) of each channel. For shifting, expression values were constitutively increased by an effect size (0 – 1.5, steps of 0.05) multiplied by the IQR. For separation, we focused on a subset of primarily bimodal channels. Using the deGate function from flowDensity (v1.42.0), negative and positive peaks were automatically gated. Expression values were subsequently reduced (negative peak) and increased (positive peak) based on the effect size (0 – 1, steps of 0.05) multiplied by IQR. To constrain the scope of the simulation and provide a realistic setting, technical effects were introduced in channels associated with each laser separately. CytoScan performance was evaluated by introducing the effects in individual files in a leave-one-out setting across the 100 Liechti_28 files.

### *In silico* simulation of biological effects

In the Liechti_28 dataset, we performed a biological perturbation experiment by removing increasing amounts of the biggest cell population in individual files. First, a balanced aggregated set of 1,000,000 cells from all files was clustered based on fluorescence parameters using FlowSOM (v2.16.0) (19) with default settings. Next, the biggest metacluster (∼40% of cells) was identified, and based on marker expression patterns, identified as CD4+ T-cells. Different amounts of CD4+ T-cells were then removed from individual files (10 – 100%, steps of 10%) by projecting the files onto the FlowSOM result and subsampling cells belonging to the metacluster. Like the technical effect simulation, output was evaluated by modifying individual files across different effect sizes in a leave-one-out setting.

## Results

### CytoScan detects anomalous FCS files in an in silico setting

We first evaluated CytoScan performance in a controlled in silico setting for a subset of samples (n=100) from a highly standardized 28-color flow cytometry cohort (Liechti_28). In this dataset, we skewed marker distributions in single FCS files (Figure 2A). The introduced effects resemble realistic technical effects: shifting, stretching and altered separation of positive and negative peak signals. As anomalies in flow cytometry data are commonly associated with parameters of specific lasers, we introduced technical effects in channels associated with each laser separately. CytoScan performance was assessed by iteratively modifying each of the Liechti_28 files (leave-one-out) and determining the percentage of correctly identified anomalous files.

**Figure 2.**
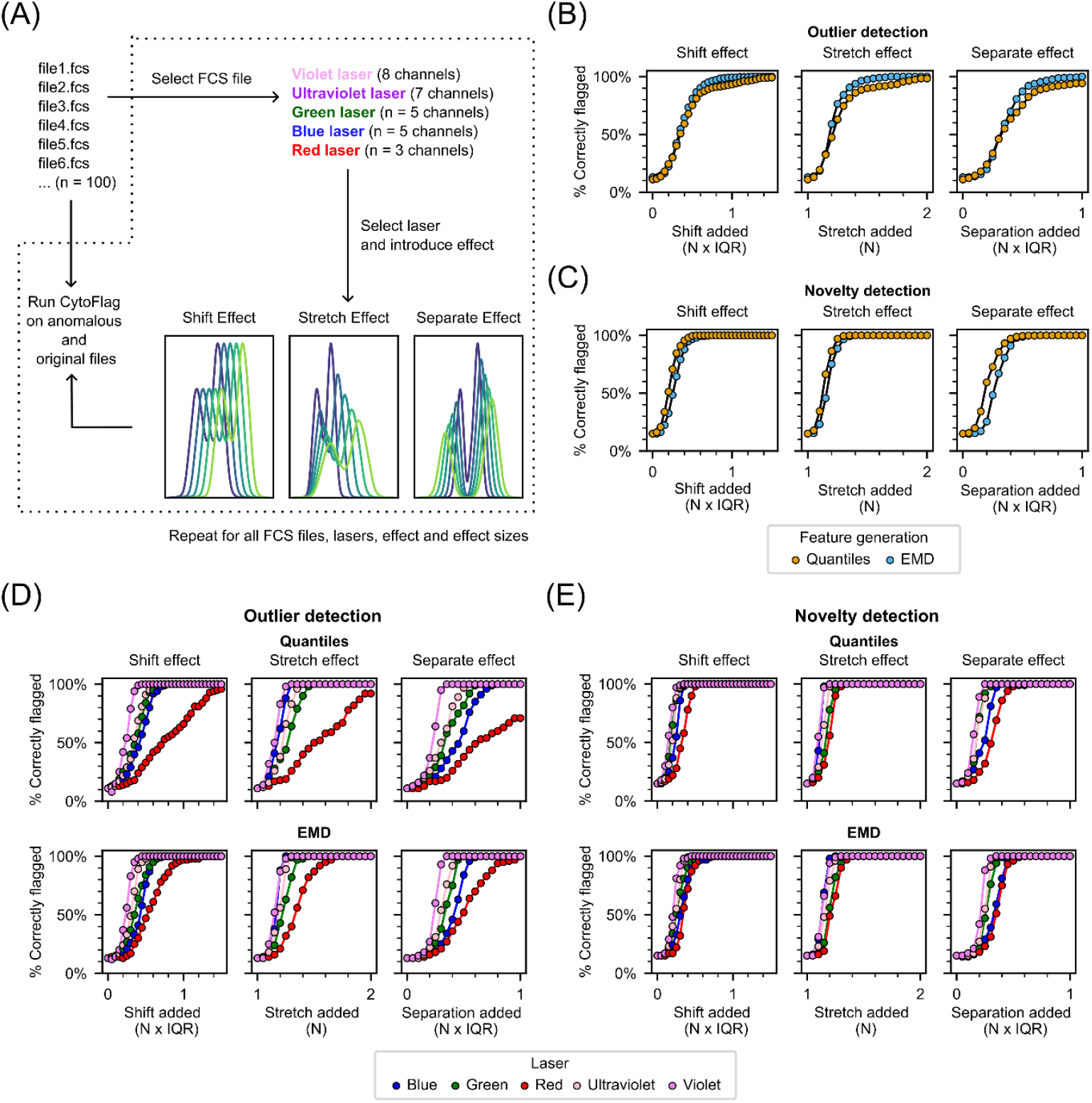
*In silico* evaluation of CytoScan detection of skewed distributions in Liechti_28 dataset. (A) Overview of in silico experimental setting. Note that the separate effect is only applied for bimodal markers. (B) Sensitivity of outlier detection for quantile and EMD-based features for shift, stretch and separate effects. Performance is shown aggregated across lasers and feature generation type. (C) Sensitivity of novelty detection for quantile and EMD-based features for shift, stretch and separate effects. Performance is shown aggregated across lasers and feature generation type. (D) CytoScan sensitivity split by laser and feature generation for outlier detection. (E) Sensitivity split by laser and feature generation for novelty detection.

CytoScan correctly identified anomalous files with different simulated technical effects, generated features, and anomaly detection algorithms (Figure 2B, C). Across all experiments, increasing the simulated effect size increased the proportion of predicted anomalies. However, results split by laser showed that effects introduced in parameters associated with the red laser (3 out of 28 parameters) are found less frequently by outlier detection, likely due to the limited number of skewed parameters (Figure 2D). Nevertheless, these results confirm that CytoScan can detect anomalies present in high-dimensional cytometry experiments if the anomalous effects are of sufficient magnitude.

In CytoScan, the novelty detection functionality leverages information from high-quality reference data to identify anomalous files. The use of such training data resulted in higher sensitivity compared to outlier detection which is performed fully unsupervised (Figure 2B,C). In particular, anomalies could be detected even when only few channels were affected, such as with the red laser (3 channels; Figure 2D,E).

Although we only recommend using CytoScan within an exploratory setting, where no samples are discarded without manual inspection, a high false positive rate may still result large numbers of anomalies that users need to validate. For the Liechti_28 dataset, we found that the false positive rate with default CytoScan settings was around

11-15% (Table S2). To evaluate false positives in a more realistic setting, we examined how CytoScan can be affected by biological variation by removing increasing amounts of the most abundant cell cluster (CD4+ T-cells, ∼40% of cells) detected by the unsupervised clustering algorithm FlowSOM (19) in the Liechti_28 dataset. Predicted anomalies were then assessed in a similar leave-one-out setting as in the previous experiment (Figure 2A). Although removal of CD4+ T-cells resulted in more predicted anomalies, for the majority of files, up to 50% of CD4+ T-cells could be removed without becoming predicted as anomalies. (Figure S2). Together, these results suggest that CytoScan can pick up (technical) anomalies in high dimensional cytometry data while remaining relatively robust to biological perturbations.

### CytoScan detects real-life technical differences

To evaluate CytoScan for real-life data, we used a location-based batch effect in a publicly available acute lymphoblastic leukemia dataset (BLN_BUE) (15) consisting of samples measured in Berlin (BLN, n=72) and Buenos Aires (BUE, n=66) using the same antibody panel (Figure 3A). First, we evaluated whether this batch effect could affect downstream analysis with FlowSOM clustering. Clustering a single BUE file together with 20 files from the BLN dataset yielded sample-specific clusters completely driven by the BUE file (Figure 3B) that could affect differential analysis of population abundance or MFIs.

**Figure 3.**
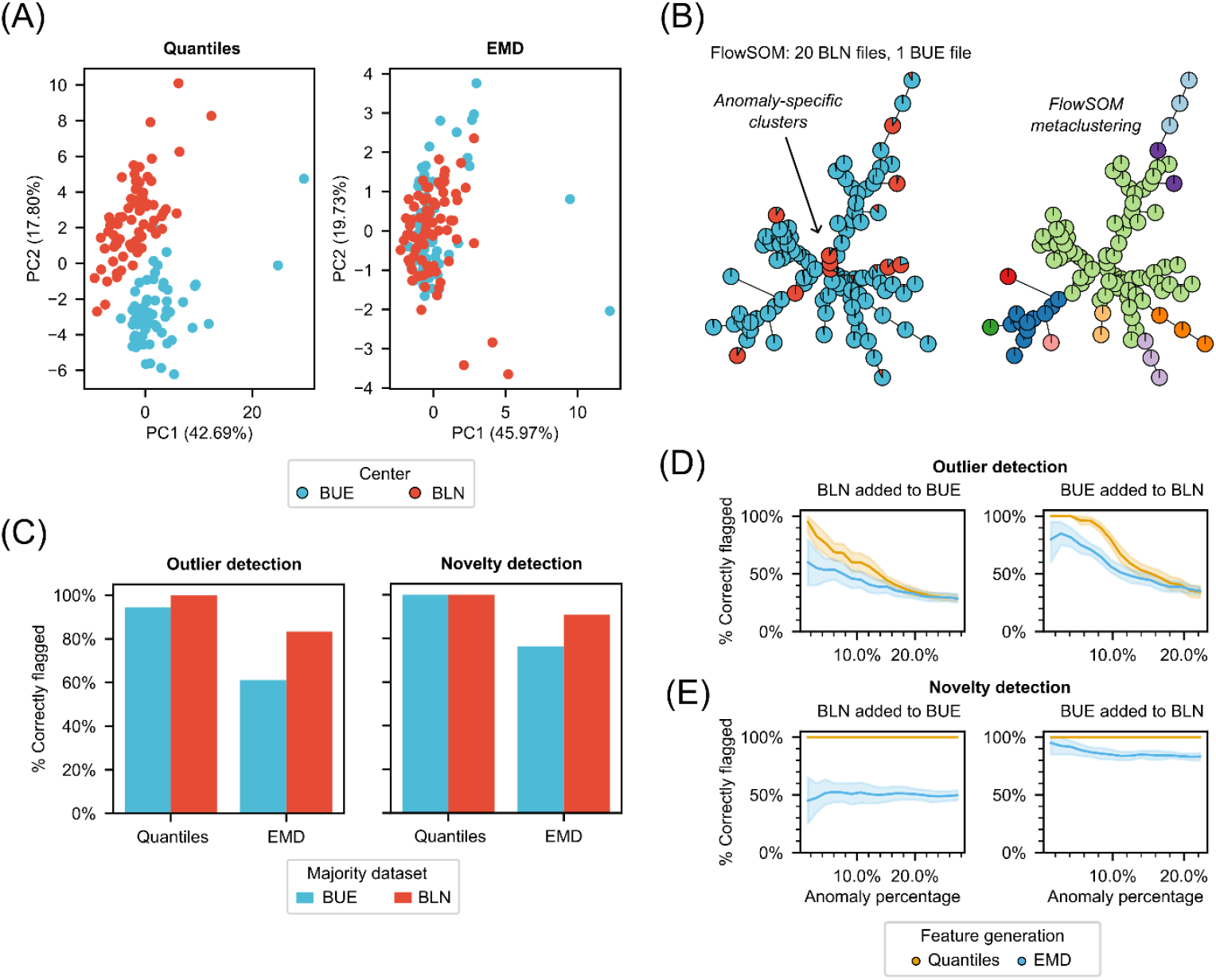
CytoScan detection of technical anomalies in real-life data. (A) Batch effect in BLN_BUE dataset visualized by principal component analysis of quantile and EMD-based features. (B) Technical differences between BLN and BUE samples drive FlowSOM clustering. Results are shown for 20 BUE samples with one added BLN sample, representing an anomalous measurement. (C) CytoScan performance in detecting single anomalies added to the full BUE and BLN cohorts. (D-E) CytoScan performance in detecting multiple anomalies added to the BUE and BLN cohorts.

Next, using this batch effect as ground truth technical anomaly, we iteratively added individual BLN samples to the complete BUE cohort (and vice-versa) and performed outlier or novelty detection using CytoScan. Quantile-based features allowed for correctly identifying 97 and 100% of anomalies when using outlier and novelty prediction, respectively (Figure 3C). EMD-based features showed lower performance (72% for outlier detection, 83% for novelty prediction; Figure 3C). This is not unexpected, as the EMD-based features already did not show a clear batch effect in the first two PCs (Figure 3A).

### Influence of batch effects on CytoScan performance

Outliers are by definition isolated events. However, technical anomalies may occur in small batches of samples that may not align with metadata used for batch effect detection. If these files share the same anomalous characteristics, they may form clusters in the generated features. Consequently, anomalous files may not be outliers anymore, preventing their detection. To evaluate the potential loss of sensitivity in cases where such batching occurs, we repurposed the experiment from the BLN_BUE dataset by adding increasingly larger sets of randomly selected files to the other dataset rather than one by one. To capture variability in the random sampling, we repeated this 20 times. As expected, sensitivity of outlier detection reduced with increasing batch size (Figure 3D) while this was not observed for novelty detection (Figure 3E). Even though novelty detection evaluates samples independently, we observed some variability for EMD-based novelty detection (Figure 3E). These variations are linked to the statistical variability of the random sampling in this simulation rather than stochasticity in CytoScan, which is prevented through setting a random seed. Taken together, we recommend that users use the anomaly detection features of CytoScan in combination with PCA and heatmap-based visualizations to rule out both anomalies and batch effects. Another option is to use a high quality reference set to use in a novelty detection setting, as this is unaffected by the relative abundance of anomalies (Figure 3E).

### CytoScan allows for extensive quality control of arbitrarily large cytometry cohorts

To demonstrate the capabilities of CytoScan in large cytometry cohorts in which manual file-by-file, marker-by-marker inspection of signals is infeasible, we evaluated measurements acquired from (primarily) bone marrow aspirates of acute myeloid leukemia (AML) patients during a large phase III clinical trial between 2014-2018 (HOVON-SAKK-132) (16). Data was acquired in a central laboratory on three different FACS CANTO II machines, providing a realistic scenario of post-hoc quality control. Experiments were performed on low-grade consumer hardware (Intel Core i3-10100T CPU, 8GB RAM). Preliminary analysis showed that CytoMDS analysis was not feasible for this dataset due to both memory requirements and excessive runtimes.

First, to assess the computational limits, we focused on the scatter parameters present in all files (FSC-H, FSC-A, SSC-H, SSC-A), resulting in a dataset (“HO132_scatter”) of 10,888 FCS files. CytoScan was able to generate features in 55 and 75 minutes for quantile and EMD-based features respectively (Figure 4A). Running outlier detection took three seconds for the quantile features and two seconds for EMD features. For novelty detection, we considered a reasonably large manually selected reference cohort (100 randomly selected files), in which case runtimes were below one second. Given that the runtimes of these steps are (approximately) linear with the number of files, these results demonstrate that CytoScan can be scaled to arbitrarily large cytometry cohorts without significant computational constraints.

**Figure 4.**
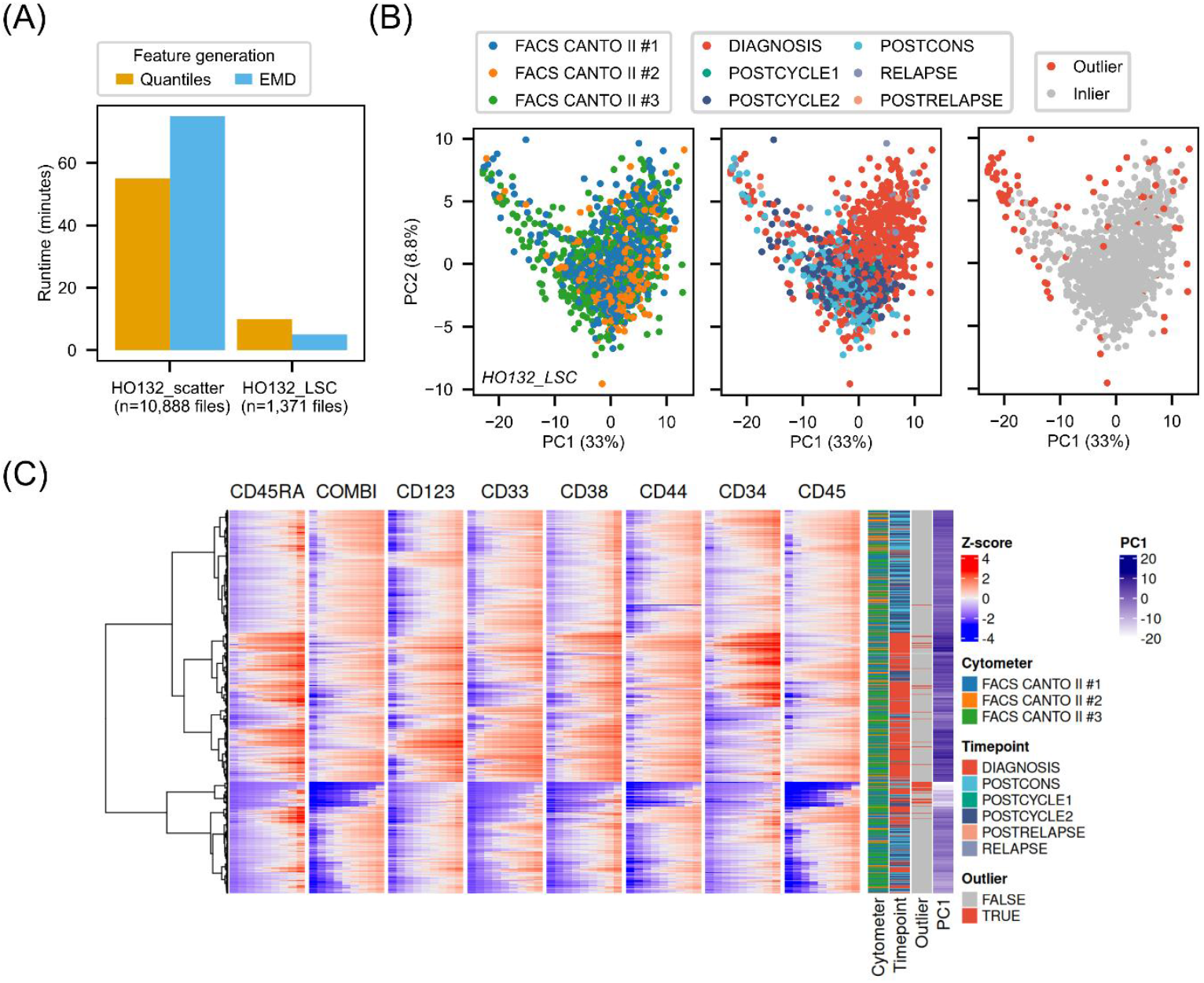
Exploratory data analysis with CytoScan in a large clinical cytometry cohort. (A) CytoScan’s runtimes for feature generation in large cohorts (HO132_scatter, HO132_LSC). Runtimes of anomaly detection (not shown) were below 5 seconds regardless of dataset. (B) Principal component analysis of quantile-based features in HO132_LSC dataset. Timepoints are along the trajectory of AML patients from diagnosis, before and after induction chemotherapy (POSTCYCLE1, POSTCYCLE2), after consolidation therapy (POSTCONS), and at or after potential relapse. (C) Heatmap of quantile-based features for each sample (rows).

Next, for a more in-depth characterization, we focused on a subset of measurements (n=1,371 FCS files) that evaluated leukemia stem cell markers (“HO132_LSC”). In this experiment, we limited the feature generation to the eight fluorescence parameters. Features were generated similarly fast (<10 minutes) (Figure 4A). PCA did not show a clear batch effect associated with the cytometer, but did show that samples clustered based on the sampling timepoint, reflecting the biology of the AML patients (Figure 4B). A clear clustering of diagnosis timepoint is expected, as in AML, the majority of cells within a sample can express a leukemic phenotype. Next, we performed outlier detection within the dataset, indicating 84 anomalous files (6.1%). In the PCA, we found an extending “tail” (PC1 low, PC2 high) consisting of a disproportionate number of outliers. To investigate this, we generated the quantile-based heatmap (Figure 4C).

This revealed a distinct group of outliers with low PC1 that shared low quantile values for CD45. Manual examination confirmed that these samples had relatively low proportions of white blood cells (WBCs), either due to large amounts of cell debris or (CD45-negative) erythrocytes.

## Discussion

Technical effects during cytometry data acquisition can lead to anomalous measurements that may lead to suboptimal and even flawed downstream analyses. We demonstrated that CytoScan can facilitate the detection of spurious technical effects and efficiently detect anomalous files. We first demonstrated this in a controlled *in silico* setting, where CytoScan could identify low proportions (1%) of anomalous files within a larger standardized high-dimensional cohort. Anomalies were still identified if they only affected few parameters in a high dimensional panel, and perturbations in T-cell subsets showed that CytoScan is relatively robust against biological variation. Next, a real-life technical effect was used to show that CytoScan can identify anomalies outside of the *in silico* setting. Lastly, we demonstrated that CytoScan scales to large cytometry cohorts, facilitating exploratory data analysis of large cytometry datasets, even in the absence of dedicated hardware.

One limitation of CytoScan is that technical effects that occur for larger batches of samples result in clustering of samples in the generated feature space, losing their outlier characteristics. Theoretically, these effects can be considered batch effects, which remains outside of the scope of CytoScan for now. Nevertheless, CytoScan users might still be interested in identifying small to medium batches of samples (5-15%) within a cohort with anomalous characteristics. Within the framework of CytoScan, these batches might still be picked up if the sensitivity of outlier detection is increased, although this also raises the false positive rate. Another solution might be to define a reference set, as this allows for novelty detection, which is performed on a per-sample basis and is therefore unaffected by groupings of samples. Alternatively, outside of the anomaly detection performed by CytoScan, we found that visualization of the features extracted by CytoScan did allow for visual detection of batch effects. Although there are batch correction algorithms available for cytometry data such as CytoNorm (2) and cyCombine (3) there remains a lack of user-friendly batch effect quantification or detection software for cytometry data.

In practice, parameter tuning of anomaly detection algorithms is difficult given that ground truth anomalies are usually unknown. As CytoScan is an exploratory tool which should be followed up by manual inspection, false negatives have more consequences than false positives (which can be excluded after manual inspection). We found that default settings resulted in 11-15% false positives in the high-dimensional Liechti_28 cohort. However, the exact rate is likely dataset-dependent and may require different hyperparameters if excessive numbers of anomalous files are infeasible to manually evaluate. In a biological perturbation experiment, CytoScan was surprisingly robust against biological fluctuations that might result in false positive classifications. This is likely because CytoScan does not take into account co-expression of markers when performing quantile or EMD-based feature generation, in contrast to a feature generation based on clustering of cells.

Ultimately, while CytoScan provides automated tools for anomaly detection, we recommend an integrated exploratory approach that includes both automated and visual identification of anomalies to reduce the labor-intensiveness of quality control. Consequentially, CytoScan helps fill an important gap in cytometry data analysis by making it easier to explore data and detect technical issues in large datasets, leading to higher-quality data and improved identification of meaningful biological patterns.

## Acknowledgements

This research was supported by a Cancer Center Amsterdam travel grant. The authors thank all participating patients and centers of the Dutch-Belgian Cooperative Trial Group for Hematology-Oncology (HOVON) and Swiss Group for Clinical Cancer Research (SAKK) 132 trial for their contribution.

## References

1. Saeys Y, Van Gassen S, Lambrecht BN. Computational flow cytometry: helping to make sense of high-dimensional immunology data. Nature Reviews Immunology. 2016;16(7):449–62.

2. Van Gassen S, Gaudilliere B, Angst MS, Saeys Y, Aghaeepour N. CytoNorm: a normalization algorithm for cytometry data. Cytometry Part A. 2020;97(3):268–78.

3. Pedersen CB, Dam SH, Barnkob MB, Leipold MD, Purroy N, Rassenti LZ, et al. cyCombine allows for robust integration of single-cell cytometry datasets within and across technologies. Nature communications. 2022;13(1):1698.

4. Hahne F, Khodabakhshi AH, Bashashati A, Wong CJ, Gascoyne RD, Weng AP, et al. Per-channel basis normalization methods for flow cytometry data. Cytometry Part A: The Journal of the International Society for Advancement of Cytometry. 2010;77(2):121–31.

5. Emmaneel A, Quintelier K, Sichien D, Rybakowska P, Marañón C, Alarcón-Riquelme ME, et al. PeacoQC: Peak-based selection of high quality cytometry data. Cytometry Part A. 2022;101(4):325–38.

6. Hahne F, LeMeur N, Brinkman RR, Ellis B, Haaland P, Sarkar D, et al. flowCore: a Bioconductor package for high throughput flow cytometry. BMC bioinformatics. 2009;10:1–8.

7. Malek M, Taghiyar MJ, Chong L, Finak G, Gottardo R, Brinkman RR. flowDensity: reproducing manual gating of flow cytometry data by automated density-based cell population identification. Bioinformatics. 2014;31(4):606–7.

8. Monaco G, Chen H, Poidinger M, Chen J, de Magalhães JP, Larbi A. flowAI: automatic and interactive anomaly discerning tools for flow cytometry data. Bioinformatics. 2016;32(16):2473–80.

9. Meskas J, Yokosawa D, Wang S, Segat GC, Brinkman RR. flowCut: an R package for automated removal of outlier events and flagging of files based on time versus fluorescence analysis. Cytometry Part A. 2023;103(1):71–81.

10. Hauchamps P, Delandre S, Temmerman ST, Lin D, Gatto L. Visual quality control with CytoMDS, a Bioconductor package for low dimensional representation of cytometry sample distances. Cytometry Part A. 2025;107(3):177–86.

11. Rogers WT, Holyst HA. FlowFP: A bioconductor package for fingerprinting flow cytometric data. Advances in bioinformatics. 2009;2009.

12. Liu FT, Ting KM, Zhou Z-H, editors. Isolation forest. 2008 eighth ieee international conference on data mining; 2008: IEEE.

13. Mocking TR, Kelder A, Reuvekamp T, Ngai LL, Rutten P, Gradowska P, et al. Computational assessment of measurable residual disease in acute myeloid leukemia using mixture models. Communications Medicine. 2024;4(1):271.

14. Liechti T, Van Gassen S, Beddall M, Ballard R, Iftikhar Y, Du R, et al. A robust pipeline for high-content, high-throughput immunophenotyping reveals age-and genetics-dependent changes in blood leukocytes. Cell Reports Methods. 2023;3(10).

15. Reiter M, Diem M, Schumich A, Maurer-Granofszky M, Karawajew L, Rossi JG, et al. Automated flow cytometric MRD assessment in childhood acute B-lymphoblastic leukemia using supervised machine learning. Cytometry Part A. 2019;95(9):966–75.

16. Löwenberg B, Pabst T, Maertens J, Gradowska P, Biemond BJ, Spertini O, et al. Addition of lenalidomide to intensive treatment in younger and middle-aged adults with newly diagnosed AML: the HOVON-SAKK-132 trial. Blood advances. 2021;5(4):1110–21.

17. Ngai LL, Hanekamp D, Janssen F, Carbaat-Ham J, Hofland MA, Fayed MME, et al. Prospective validation of the prognostic relevance of CD34+ CD38–AML stem cell frequency in the HOVON-SAKK132 trial. Blood, The Journal of the American Society of Hematology. 2023;141(21):2657–61.

18. Zeijlemaker W, Kelder A, Oussoren-Brockhoff Y, Scholten W, Snel A, Veldhuizen D, et al. A simple one-tube assay for immunophenotypical quantification of leukemic stem cells in acute myeloid leukemia. Leukemia. 2016;30(2):439–46.

19. Van Gassen S, Callebaut B, Van Helden MJ, Lambrecht BN, Demeester P, Dhaene T, et al. FlowSOM: Using self-organizing maps for visualization and interpretation of cytometry data. Cytometry Part A. 2015;87(7):636–45

