## Supplementary Information for "CytoScan: Automated detection of technical anomalies for cytometry quality control"

**Table S1. Feature generation methods in CytoScan (v1.0.0)**

| Method | Description | Arguments | # Features | Requires aggregate |
|---|---|---|---|---|
| Summary statistics | Expression value at mean, median. Expression range of standard deviation, interquartile range. | None | 4 x $N_{channels}$ | No |
| Binning | Proportion of cells within each bin. | Number of bins | $N_{bins}$ x $N_{channels}$ | Yes* |
| flowFP fingerprint | Proportion of cells within each flowFP bin. | Number of recursions | $2^{Nrecursions}$ | Yes |
| Quantiles | Expression value at quantile. | Number of quantiles | $N_{quantiles}$ x $N_{channels}$ | No |
| Earth mover's distance | Distance with respect to aggregate distribution. | None | $N_{channels}$ | Yes |

Note: Aggregate is used to set determine bin boundaries. While binning can theoretically be performed on single files similarly as quantiles, this could result in shifted files having identical features.

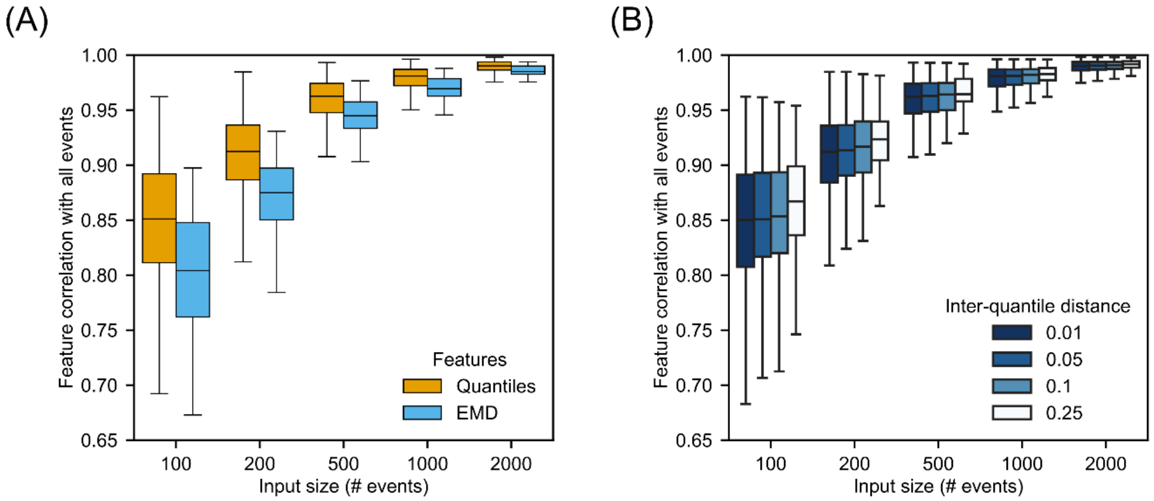


**Figure S1. Concordance of features generated in downsampled and full FCS files.** (**A**) EMD versus quantile-based (0.1 interquantile distance) feature generation. (**B**) Effects of interquantile distance and concordance of downsampled features.

**Table S2. False positive rate (FPR) of CytoScan in the Liechti_28 dataset.**

| Feature generation | Flagging method | FPR |
|---|---|---|
| **Quantile** | Outlier | 11% |
| **EMD** | Outlier | 13% |
| **Quantile** | Novelty | 15% |
| **EMD** | Novelty | 15% |

The FPR of novelty detection was evaluated in a leave-one-out setting. The FPR is dataset-dependent. If users observe many incorrectly flagged files, consider hyperparameter tuning.

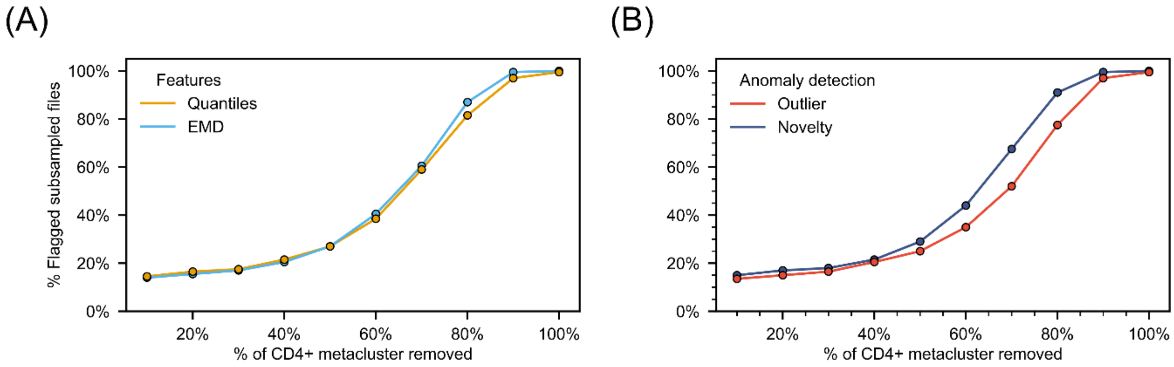


**Figure S2. Sensitivity to biological changes.** (**A**) Percentage of detected anomalies for quantile and EMD-based features across a range of biological perturbations.(**B**) Percentage of detected anomalies for outlier and novelty detection across a range of biological perturbations.
